# GDF15 modifies NAD+ salvage pathway and dampens immune response in mitochondrial disease

**DOI:** 10.64898/2026.10.31.763316

**Authors:** Anastasiia Marmyleva, Nahid Khan, Liliya Euro, Sonja Jansson, Anu Suomalainen

## Abstract

Mitochondrial integrated stress response (ISRmt) is a main mammalian metabolic stress pathway, with local ISRmt and a systemic response via circulating metabokines GDF15 and FGF21 as early signs of the response in mitochondrial diseases. FGF21 has been reported to regulate ISRmt progression and nutrient homeostasis, but the roles of GDF15 have remained unclear. Here, we explored the effects of inactivation of FGF21 or GDF15 or them together in Deletor mice. These mice endogenously activate the full ISRmt and metabokines as part of their mitochondrial muscle disease. GDF15 inactivity upregulated the skeletal muscle enzyme NMRK2, which converts nicotinamide riboside (NR) to nicotinamide mononucleotide (NMN) and NAD+. FGF21 was required for glutathione homeostasis. GDF15 inactivation also induced immune cell invasion in the muscle, aggravated by FGF21 loss, suggesting synergistic anti-inflammatory effects. Our findings indicate that in addition to their systemic effects, FGF21 and GDF15 are autocrine controllers of redox homeostasis in mitochondrial disease muscle.

**Highlights:**

- GDF15 and FGF21 are autocrine redox regulators in mitochondrial myopathy
- GDF15 remodels NAD salvage pathway via NRMK2 in the skeletal muscle
- FGF21 is necessary for glutathione synthesis in the skeletal muscle
- GDF15 and FGF21 synergize as anti-inflammatory in mitochondrial muscle disease

## Introduction

Systemic metabolic signaling by molecules that are secreted from one tissue and affect the whole organismal metabolism is increasingly appreciated. Such factors can also be messengers of disease, tuning the organism to face the endogenous stress. In mitochondrial diseases, the growth differentiation factor 15 (GDF15) and fibroblast growth factor 21 (FGF21) are such signalers, conserved in mouse models and human patients^1,2^. Their specificity and sensitivity to the disease led to their validation as serum biomarkers, used in neuromuscular and mitochondrial clinics and as readouts in clinical trials^2–6^. Despite their clinical use, the physiological relevance of GDF15 and the potential crosstalk of the two metabokines in the context of mitochondrial diseases are still unclear.

Both FGF21 and GDF15 were originally identified in the mitochondrial muscle disease model, the “deletor” mice, and then found to be similarly induced in human patients with diseases involving mitochondrial DNA (mtDNA) instability or mtDNA expression defect^3,5,7,8^. GDF15 and FGF21 are highly expressed already in the early stage of the disease process, as the initial step of mitochondrial integrated stress response, ISRmt^9^. Progression of ISRmt is partially FGF21 dependent: if FGF21 is inactivated in mice with mitochondrial muscle disease, the second stage components of ISRmt (glucose uptake to muscle, adipose tissue, heart and dorsal hippocampus, *de novo* serine biosynthesis, transsulfuration)^10,11^, remain uninduced^9^. FGF21 appears therefore to be a major regulator of ISRmt progression in chronic disease.

The tissue-specific and hormone-like functions of GDF15 have garnered significant attention since its discovery over 25 years ago^12^. GDF15 has been reported to suppress appetite via central nervous system mediated mechanism, linked with pregnancy-related hyperemesis, cachexia and anorexia^13,14^. GDF15 has also anti-inflammatory functions, which may partially contribute to its protective effects in obesity as it reduces inflammation in adipose tissue^15–17^. Clinical trials of GDF15 inhibition in cancer-related anorexia/cachexia are ongoing in phase II trials with promising results^18^. Whether chronically elevated GDF15 has protective, neutral or deleterious effects in mitochondrial diseases awaits pathomechanistic knowledge.

Here, we report the physiological roles of GDF15 and FGF21 independently and together in mitochondrial myopathy, using the deletor mice that endogenously express these metabokines in their skeletal muscle and heart. These mice carry a 39 bp insertion of *TWNK*-gene, encoding Twinkle, the nuclear-encoded replicative helicase of mtDNA. The mutation is homologous to that in mitochondrial myopathy patients and affects the linker region of the hexameric helicase. Deletor mice manifest all hallmarks of mitochondrial myopathy in their muscle after 18 months of age^1,7,9,10,19–21^. Deletors also were the basis to provide preclinical data for biomarker and therapy development for the disease^3,5,8^. Here we used a GDF15-neutralizing antibody, which blocks the metabokine binding to GFRAL receptor and thereby impedes its endocrine function^22^. We report GDF15 engagement in NAD+ metabolism, especially that of NAD+ precursor nicotinamide mononucleotide (NMN) in skeletal muscle, but not in other tissues. Furthermore, GDF15 prevents activation of immune system in response to mitochondrial DNA replication deficiency. Part of these effects also require FGF21, indicating the synergistic and specialized functions of the two metabokines, forming an important metabolic signaling system.

## Results

### GDF15 inhibition does not impact morphological hallmarks of mitochondrial disease

The phenotype and molecular responses of mitochondrial myopathy in deletors (DEL) were characterized previously in detail^1,5,7,9,10^. Starting from 14 months of age, these mice start to increasingly express both FGF21 and GDF15.

Here, we utilized 2-year-old deletors and deletor-FGF21 knockout (DELFKO) mice^9^, subjected also to inhibition of by a GDF15-neutralizing antibody (INH), injected once per week for one month in mice that manifested robust mitochondrial myopathy **(Fig. 1A)**. Blockade of GDF15 signaling did not compromise the overall well-being of mice during the four weeks of treatment, when assessed by weight, food consumption and glucose tolerance **(Fig. S1A,B,C)**. On the systemic level, loss of FGF21 caused metabolic switch from glucose to lipid utilization as the preferred cellular energy source by the muscle with mitochondrial disease. Loss of GDF15 activity in DELFKO mice shows a tendency to decrease the respiratory exchange ratio **(Fig. 1B,C)**.

**Figure 1.**
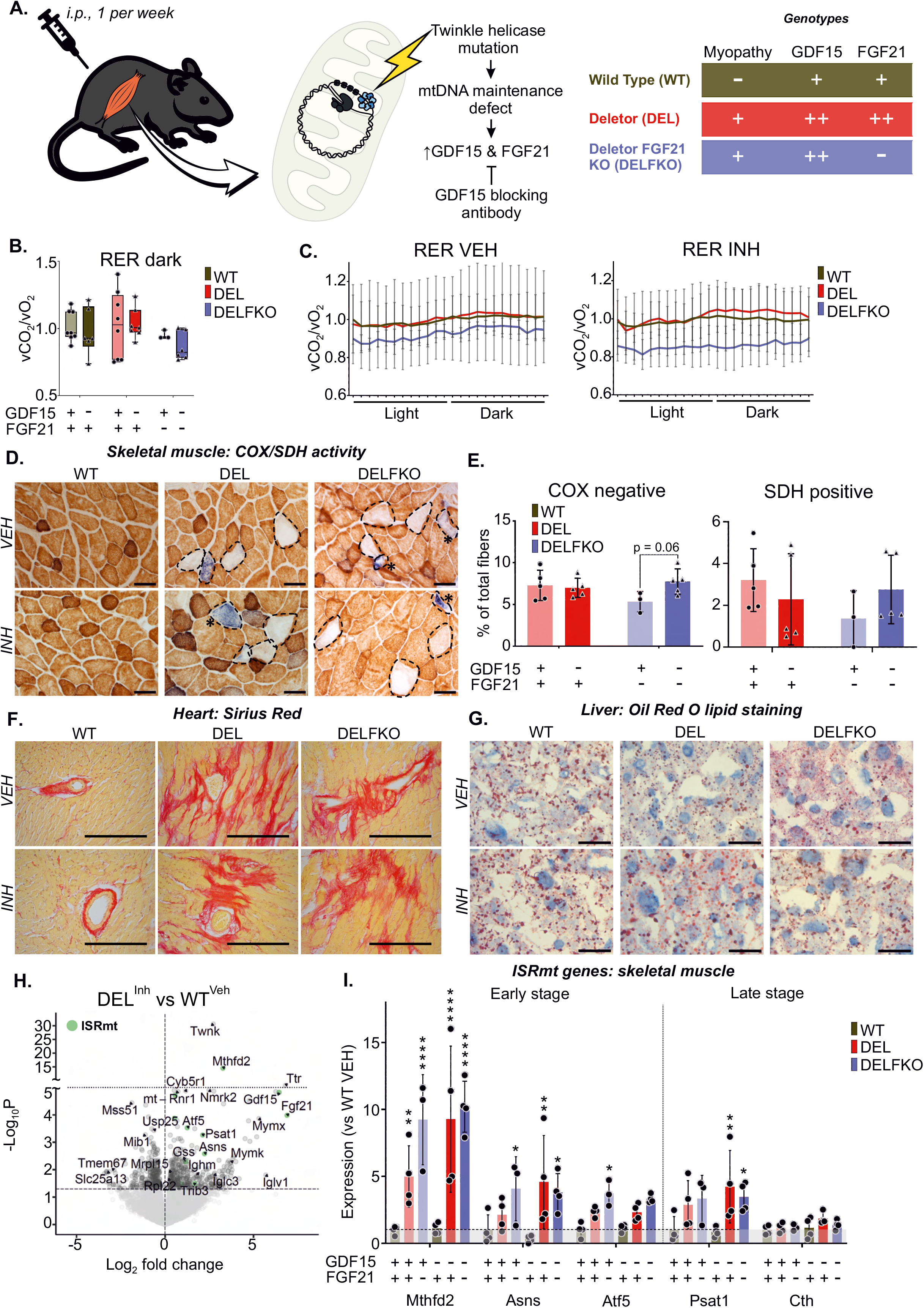
GDF15 inhibition does not worsen major hallmarks of the disease. (A) An experimental layout of GDF15 inhibitor treatments. (B) Respiratory exchange ratio (RER), CLAMS metabolic cages. n = 3-8/group. (C) RER curves plotted separately for vehicle and inhibitor treatments. n = 3-8/group. (D) COX/SDH histochemical activity analyses on frozen skeletal muscles to detect activities of mitochondrial complexes IV (COX, cytochrome c oxidase) and II (SDH, succinate dehydrogenase). Brown – COX, blue – SDH positive. Scale bar for 80 um; n = 3-6/group. (E) Quantification of COX negative and SDH positive fibers from Fig. 1D. Average number of fibers quantified = 1670; n = 3-5/group. (F) Fibrosis, cardiac muscle. Sirius Red staining, (red) in the heart. Scale bar for 200 um; n = 3- 6/group. (G) Lipid content in mouse liver, frozen sections. Oil Red O and hematoxylin staining. Scale bar for 80 um; n = 3-6/group. (H) Differentially expressed genes, deletor mouse skeletal muscle. Volcano plot with inhibition of GDF15 signaling as compared to vehicle treated WT. n=4. (I) ISRmt gene expression in skeletal muscles. n=3-4/group. Abbreviations: WT – wild type, DEL – deletor, DELFKO – deletor FGF21 knockout, VEH – vehicle, INH – inhibitor.

To test the effects of GDF15 and FGF21 for mitochondrial disease hallmarks, we analyzed the amount of mitochondrial DNA (mtDNA) and its deletions, as well as histochemical activity of respiratory chain enzymes cytochrome c oxidase (COX, partially encoded by mtDNA) and succinate dehydrogenase (SDH, nuclear encoded) *in situ* on frozen sample of skeletal muscle of the mouse lines. Although depletion of FGF21 in deletors mildly decreased the total mtDNA amounts, the effect was neither aggravated nor rescued by GDF15 inhibition **(Fig. S1D)**. Additionally, neither GDF15 nor FGF21 affected the mtDNA deletion load in mouse skeletal muscle **(Fig. S1E,F)**. The quadriceps femoris (QF) muscles of all mice with myopathy background (DEL and DELFKO) displayed COX-negative and SDH-positive fibers **(Fig. 1D,E).**

The absence or presence of FGF21 or GDF15 separately did not affect the abundance of the disease-related COX-/SDH+ muscle fibers. Inhibition of both metabokines (DELFKO plus GDF15 antibody) mildly increased the amount of COX-negative fibers **(Fig. 1E)**.

Previous reports show that the skeletal muscle is the primary affected organs in deletor mice, the heart does not show functional defects but does show changes in metabolism, and the liver displays secondary lipid metabolic changes in response to the disease^1,7,9,10^. Despite no functional difference in the hearts, we detected large fibrosis areas, which were present in mice with deletor background (DEL and DELFKO) but independently of either metabokine **(Fig. 1F)**. In the presence of GDF15 and FGF21, the liver tissue of Deletor mice shows a decreased amount of both intra- and extracellular lipid droplets **(Fig. 1G)**. FGF21KO restored the extracellular amount of lipid droplets in Deletors to the wild type level, and GDF15 inhibition normalized the intracellular lipid droplets to the wildtype level, indicating a compartment-specific effect on lipid storage of both metabokines **(Fig. 1G).**

Induction of the mitochondrial integrated stress response is a molecular hallmark of mitochondrial myopathy, also upregulated in the Deletor skeletal muscle. The transcriptomic components of ISRmt included induction of FGF21 and GDF15, M*THFD2, ATF5* and *ASNS* of the first-stage ISRmt and *PSAT1, PHGDH, CBS* and *CTH* of the 2^nd^ stage ISRmt. Loss of FGF21 increased *MTHFD2* and *ATF5* transcription further, while *ASNS* induction was dampened, indicating local effects of FGF21 in the skeletal muscle. GDF15 had no remarkable effect on these **(Fig. 1H,I; S1G)**. Second stage components of ISRmt were mildly dampened by FGF21 KO but showed no dependence on GDF15. EIF2alpha and mTORC1 activation, part of second stage response, were not affected by either metabokines **(Fig. 1SH,I,J)**. The heart also shows induction of ISRmt components, whose levels were regulated by FGF21 but not GDF15 **(Fig. S1K)**. On the other hand, the liver did not activate the stress response components except for the *ASNS*, which was strongly induced in deletors. This effect was exacerbated by FGF21 deletion but dampened by GDF15 inhibition **(Fig. S1L)**.

Taken together, these data indicate that GDF15 inhibition is well-tolerated in skeletal muscle, the main manifesting organ in mitochondrial myopathy. GDF15 and FGF21 show a synergistic role in systemic and liver lipid metabolism, mobilization of lipids from tissue and usage as a fuel. FGF21 has a special role in the regulation of ISRmt response in the skeletal muscle, with no contribution by GDF15.

#### GDF15 and FGF21 have separate roles in redox metabolism

Redox balance in cells is guarded by reduced and oxidized NAD and glutathione forms. The deletor heart and muscle are FGF21-dependent for transsulfuration pathway induction, and glutathione, the end- product of transsulfuration, is the main direction of glucose carbon flux in these tissues^10^. These findings prompted us to explore the redox balance of deletor mice in the presence or absence of FGF21 and GDF15 activity. The deletor skeletal muscle had low GSH (reduced glutathione)/GSSG (oxidized) ratio in all mice lacking FGF21, which together with the findings of FGF21 regulation of transsulfuration pathway, pointed to decreased production of GSH **(Fig. 2A)**. GDF15 presence or absence did not affect GSH/GSSG ratio.

**Figure 2.**
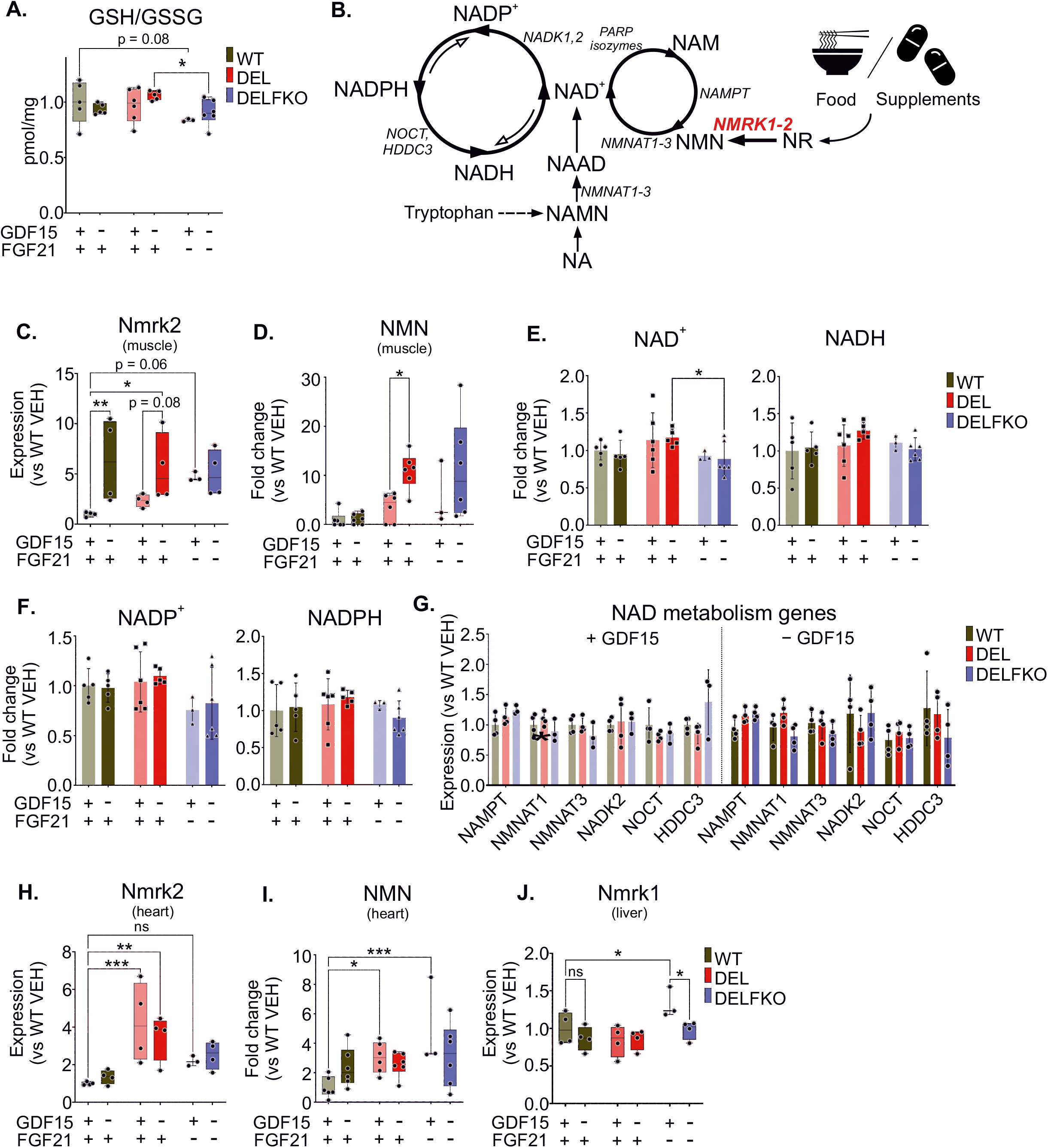
GDF15 alters cellular redox balance by directing glutathione metabolism and NAD biosynthesis. (A) Reduced (GSH) and oxidized (GSSG) forms of glutathione in skeletal muscle. n = 3-6/group. (B) NAD synthesis; nicotinamide riboside (NR), schematic representation. (C) Nmrk2 gene expression in skeletal muscle. n = 3-6/group. (D) Nicotine mononucleotide amount (NMN) in skeletal muscle. n = 3-6/group. (E-F) NAD metabolite amounts in skeletal muscles. n = 3-6/group. (G) NAD synthesis gene expression, n=3-4. (H) Nmrk2 gene expression in the heart. n = 3-6/group. (I) NMN amount in the heart. n = 3-6/group. (J) Nmrk1 gene expression in the liver. n = 3-6/group. Abbreviations: WT – wild type, DEL – deletor, DELFKO – deletor FGF21 knockout, VEH – vehicle, INH – inhibitor. NMRK1 and 2 – Nicotinamide Riboside Kinase 1 and 2, Star significance: ns – nonsignificant, p>0.05, * - p ≤ 0.05, ** - p ≤ 0.01, *** - p ≤ 0.001, **** - p ≤ 0.0001.

However, the absence of GDF15 activity had a role in the second main redox active system, the NAD+ synthesis. In this pathway, nutrient-derived vitamin B3 forms are metabolized to NAD+. Nicotinamide riboside (NR) is one of these NAD+ precursors, a nutrient-derived vitamin B3 form. First, NR is converted into nicotinamide mononucleotide (NMN) by NR kinases, and NMN enters the salvage pathway to form NAD+ **(Fig. 2B)**. In the skeletal muscle and the heart, the kinase is NMRK2, and in the liver NRMK1. We found that inhibition of GDF15 activity activated NMRK2 expression up to 10-fold in the skeletal muscle of both control and deletor mice **(Fig. 2C)**. In line with the increased NMRK2 activity, GDF15 inhibition resulted in up to 10-fold increase of muscle NMN **(Fig. 2D)**. NMN changes occurred only in the context of mitochondrial myopathy. In the control animals, NMRK2 similarly increased in the absence of GDF15, but this did not change the steady-state NMN amount **(Fig. 2C,D).** FGF21 absence only mildly lowered NAD+ levels in deletors. On the other hand, GDF15 did not affect NADH, NADP, or NADPH levels of the muscle, and the expression of the NAD-cycling enzymes remained stable **(Fig. 2E,F,G)**.

In the deletor heart, NMRK2 was induced 4-fold, but was not affected by either metabokine **(Fig. 2H).** Similarly, heart NMN levels mildly increased in deletor background **(Fig. 2I)**. In the liver, FGF21 presence suppressed NMRK1 whilst GDF15 presence managed to normalize its levels to wildtype level **(Fig. 2J)**.

These results indicate that FGF21 and GDF15 remodel redox metabolism, with specific downstream effects: FGF21 controls glutathione homeostasis, while GDF15 effect is on NAD-salvage pathway in the context of mitochondrial myopathy. Importantly, the effects were tissue specific.

#### GDF15 loss activates immune system in MM mice

The RNA-sequencing indicated that GDF15 inhibition activated a wide-spread inflammatory signal in the skeletal muscle of both the deletors and controls, with mRNAs of B-cell and plasma cell activation (Ighm - immunoglobulin heavy constant Mu; Igkc -immunoglobulin kappa constant), b-cell receptors (Cd79b, MS4A1), complement system activation and inflammatory leukotriene synthesis **(Fig. 3A, S2A)**. The effect was the most pronounced in the presence of FGF21 when GDF15 was inhibited **(Fig. 3B,C; S2A)**. The histological analysis of skeletal muscle of Deletors and controls showed clear signs of inflammation. Both CD45, a marker for nucleated hematopoietic cells (B and T cells), and CD68, the marker of macrophages and monocytes, presented large clusters in muscles of mice lacking GDF15 activity. This effect was lowered by removal of the FGF21 signal although there was still stronger immune cell infiltration in these muscles compared to vehicle-treated controls **(Fig. 3D)**. These observations reflect our transcriptomics findings.

**Figure 3.**
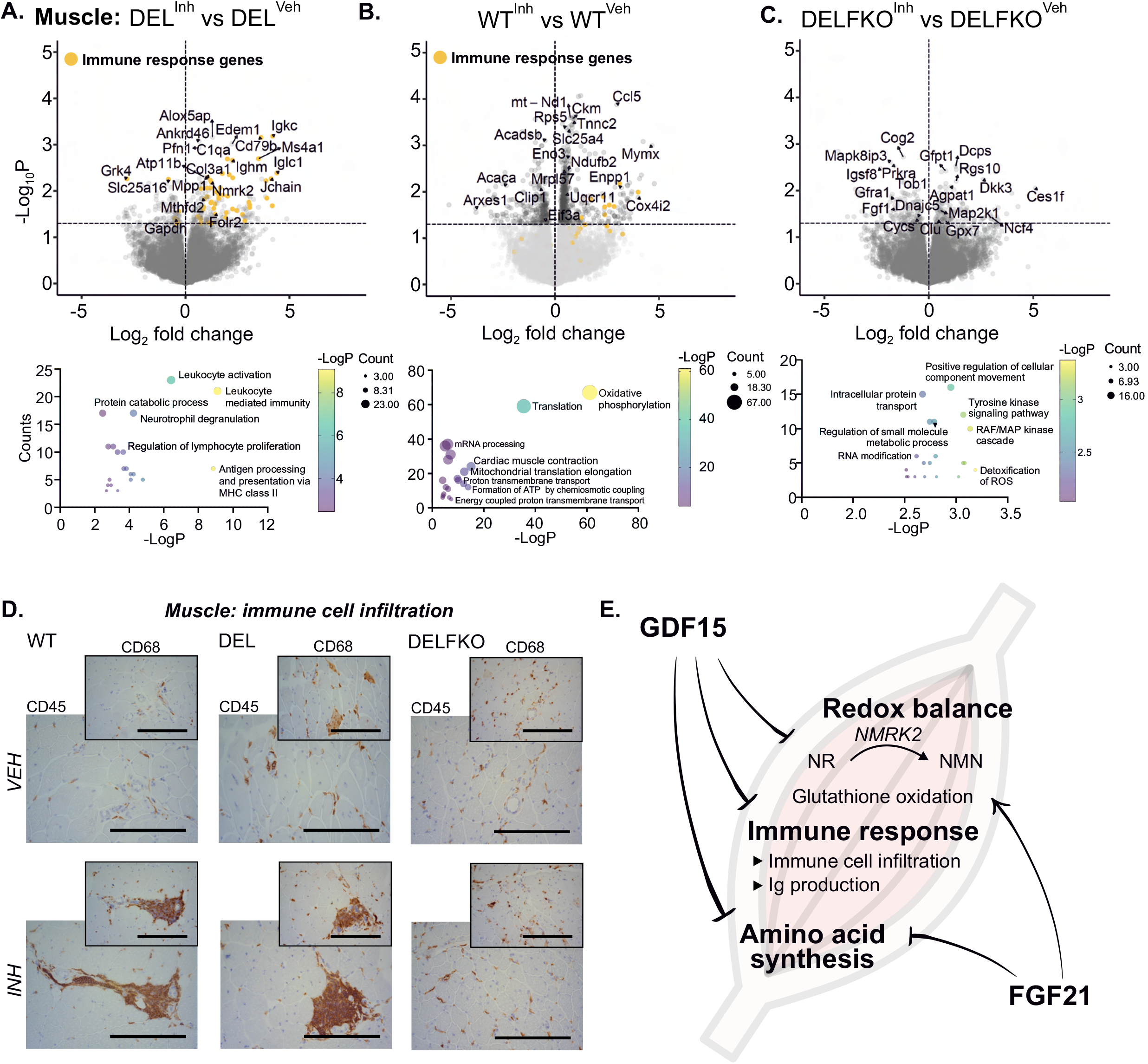
GDF15 suppresses immune activation in MM muscles. (A-C) Differentially expressed genes after inhibited GDF15 activity in (A) deletors, (B) wild types and (C) deletors without FGF21; volcano plots and pathway analyses; n=4. (D) Plasma cells (CD45) and macrophages (CD68) in skeletal muscles; immunohistochemical staining. Scale bar for 200 um; n = 3-6/group. (E) Schematic summary of the main functions of GDF15 and FGF21 in muscle tissue. Abbreviations: WT – wild type; DEL – deletor; DELFKO – deletor FGF21 knockout; VEH – vehicle; INH – inhibitor. Star significance: ns – nonsignificant, p>0.05, * - p ≤ 0.05, ** - p ≤ 0.01, *** - p ≤ 0.001, **** - p ≤ 0.0001. n=3-6/group.

While induced mRNAs suggested that GDF15 had a singular role in inflammation suppression, the presence of FGF21 when GDF15 was inhibited showed severe inflammatory changes, suggesting that without GDF15, FGF21 is inflammatory **(Fig. 3E)**. Overall, our findings support the role of GDF15 as a suppressor of inflammation in the context of skeletal muscle and mitochondrial myopathy.

### Metabokine-specific metabolome remodeling

We then explored the effects of the two metabokines on skeletal muscle and heart metabolomics. Changes of the metabolome in both organs were most profound under combined signaling inactivation of FGF21 and GDF15 in deletor mice.

The deletor mouse muscle shows an increase in amino acids, such as glutamine, glutamate, and lysine. Free amino acid levels increased by FGF21KO and further by GDF15 inhibition. When both metabokines were inhibited, charged amino acids (such as Arg, Lys, Asp, and Glu) were further increased, as well as Ser and Gly, which are part of serine-glycine-one carbon cycle response to mitochondrial disease (**Fig. 4A-E)**. Palmitoleic acid, a monounsaturated medium-chain omega 7 fatty acid (16:1n-7), also presented similar dependance on both metabokines. FGF21 strongly inhibited its steady-state levels exemplified by a 4-fold increment in its levels after FGF21 depletion, while GDF15 inhibition restores it to near wild type level **(Fig. S3A).** Wild type muscle showed little changes after GDF15 inhibition **(Fig. S3B)**.

**Figure 4.**
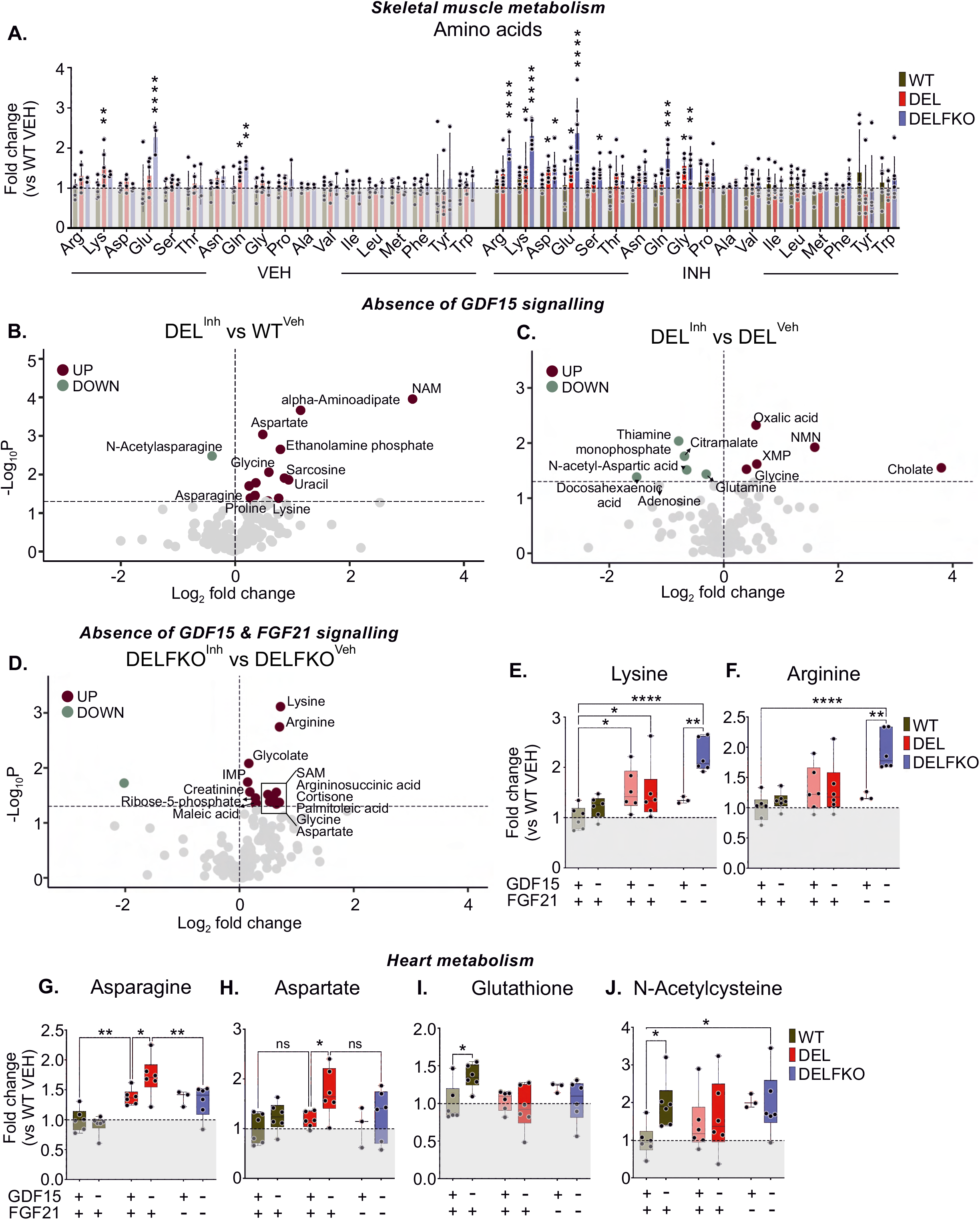
GDF15 and FGF21 absence further disbalances amino acid metabolism. (A) Fold changes in amino acid levels of skeletal muscles. (B-C) Significantly changed metabolites in muscles of deletor mice with inhibited GDF15 activity as compared to vehicle-treated (B) healthy WT mice or (C) mice of the same genotype. (D) Significantly changed metabolites in muscles deletor mice lacking both GDF15 and FGF21 activities. (E-F) Metabolic levels of (E) Lysine and (F) Arginine. n = 3-6/group. (G-H) Metabolic levels of (G) Asparagine, (H) Aspartate, (I) total Glutathione and (J) N-Acetylcysteine in the heart. n = 3-6/group. Abbreviations: WT – wild type; DEL – deletor; DELFKO – deletor FGF21 knockout; VEH – vehicle; INH – inhibitor; NAM – nicotinamide; NMN – nicotinamide mononucleotide; XMP - xanthosine monophosphate; SAM - S-adenosylmethionine; IMP – Inosine monophosphate. Star significance: ns – nonsignificant, p>0.05, * - p ≤ 0.05, ** - p ≤ 0.01, *** - p ≤ 0.001, **** - p ≤ 0.0001. n=3-6/group.

In the heart of deletor mice, the absence of GDF15 activity also changed free amino acids, especially increasing aspartate and lowering asparagine **(Fig. 4G,H; S4C).** Aspartate is made in mitochondria and used in asparagine synthesis by asparagine synthetase (ASNS), an early responder in ISRmt in deletors. The increase of ASNS expression did not depend on GDF15’s presence, but the induction was partially dampened when FGF21 was absent; FGF21 is also part of early ISRmt **(Fig. S4D)**. FGF21 alone also affected the redox metabolism as its absence led to increased levels of cystathionine and N- acetylcysteine, intermediates of glutathione synthesis **(Fig. S3E,F)**.

When both metabokine activities were absent, the deletor heart showed few changes, and unlike the skeletal muscle, had no increases of amino acids. The changes included a decrease in some tricarboxylic cycle metabolites (fumarate, malate) as well as carnosine, glyoxylic acid, and 4-imidazole acetic acid that are connected with each other by histidine/histamine metabolism **(Fig. S3G)**. In wild type mouse hearts, GDF15 inhibition led to high steady-state N-acetyl-L-cysteine and total glutathione levels, both being part of transsulfuration pathway and important for redox balance in the heart **(Fig. 4I,J; S3H)**.

All in all, our metabolomics data support the synergistic roles of GDF15 and FGF21 in particular for amino acid metabolism.

### GDF15 exerts organellar effects that are genotype- and tissue-specific

In skeletal muscle of healthy WT mice, GDF15 affected expression of genes specific to mitochondrial electron transport chain, correlating negatively with mitochondrial respiratory chain components, translation (both mitochondrial and cytoplasmic), and contraction machinery **(Fig. S4A,B)**. In hearts of DELFKO mice, these essential programs were also inhibited by GDF15 **(Fig. S4C)** to a higher extent than in WT and DEL hearts upon GDF15 inhibition **(Fig. S4D,E)**. On the other hand, transcriptional changes in livers did not replicate similar changes as in the muscle or heart **(Fig. S4F-H)**. Endoplasmic reticulum (ER) and Golgi-associated pathways were the most GDF15-sensitive pathways in livers of deletor mice with FGF21 (DEL; **Fig. S4F**).

The data from different tissues further highlights the tissue-specific sensitivity to GDF15. Moreover, mitochondrial dysfunction and loss of FGF21 increases the tissue susceptibility to GDF15 effects on organellar functions.

## Discussion

We report here the separate and combined effects of two metabokines, GDF15 and FGF21, in mice with mitochondrial muscle disease and in aged healthy mice **(Fig. 3E)**. The deletor mouse, a model for mitochondrial myopathy, has played a key role in identification of both FGF21 and GDF15 in mitochondrial diseases^1,2^ as the findings were then shown to apply also in human patients and developed as serum biomarkers for mitochondrial disease ^2–4,6^. The mice upregulate endogenously both metabokines as part of ISRmt as the early, subclinical stage of the disease^1,2,4,5^. Here, when independently and together depleting both metabokine activities in the context of mitochondrial disease and healthy controls, we discovered their interesting novel roles of these metabokines in redox metabolism. Lack of GDF15 in the context of mitochondrial disease highly upregulated NMRK2, the enzyme that converts NR to NMN. Consequently, steady-state NMN levels increased, suggesting that GDF15 activity regulates how efficiently nicotinamide riboside is converted to NMN in the muscle. This finding was limited to skeletal muscle and not present for example in the heart. The chronic upregulation of GDF15 could also contribute to NAD depletion, because of inefficient metabolism of NR to NMN. This would be consistent with the NAD depletion reports in skeletal muscles of mitochondrial myopathy patients^8^. The findings also emphasize that the contribution of different B3 vitamin forms (niacin, NR, NMN, nicotinamide) to NAD pool may vary in across tissues; important to acknowledge as these NAD-precursors are used in large doses in mitochondrial diseases^8^ and treatment trials for degenerative diseases.

The glutathione redox system is the other main redox regulator of cellular homeostasis. We found that the skeletal muscle of deletor mice the loss of FGF21 lowered the glutathione pools. Our results and previous data support the conclusion that the activation of transsulfuration pathway, regulated by presence of FGF21^9^, maintains the reduced state via GSH pool in the skeletal muscle in mitochondrial muscle disease. However, in the heart of healthy animals, the loss of GDF15 activity, but not FGF21, increased the steady-state N-acetyl-L-cysteine and the total glutathione pool. The data indicate that the metabokines have tissue and context specific regulatory roles on redox-status in health and disease.

We found the third major effect of GDF15 and FGF21 to be on inflammation. The presence of FGF21 and GDF15 suppressed inflammatory changes in deletor skeletal muscle. When both metabokines were inactive, the muscle showed degenerative changes as well as clusters of immune cells. These clusters included both plasma cells, indicating muscle-derived antigen presentation and immune activation, as well as macrophages pointing to muscle damage. The contribution of redox state on inflammation may be a factor. Knockout mouse models for mitochondrial proteins were reported to trigger immune responses^23,24^, and those with mtDNA maintenance defects also to modulate immune activation^25^. Our data indicate that induction of ISRmt with its early responders to mitochondrial myopathy – FGF21 and especially GDF15 – are anti-inflammatory in the skeletal muscle already in the subclinical diseases stage.

Many of the previous studies exploring GDF15 functions were conducted on mice that received recombinant GDF15 or were knockouts of GDF15^26–29^. Our data on endogenous regulation offered an excellent possibility to explore the interaction of GDF15 and FGF21 in deletor context that has previously been relevant for patient diagnosis and also treatment^6,8^. One of the main reported GDF15 roles is loss of appetite and weight^26,30–32^. We did not find such effects during the intervention, even though the remodeling of the redox state and amino acid metabolism was remarkable, especially in the context of mitochondrial disease.

Overall, the current and previous data show that FGF21 prevails as a dominant regulator of ISRmt progression, regulating glucose uptake and redirecting carbon flux toward folate cycle–dependent pathways, including *de novo* serine biosynthesis, purine nucleotide synthesis, and the transsulfuration pathway driving cysteine and glutathione production^3,6,9^. GDF15, in turn, has important roles in tissue- specific NAD metabolism and inflammation. Together, these metabokines appear to shape the adaptive response to mitochondrial myopathy by modulating systemic metabolism while simultaneously restraining excessive local immune activation. In healthy animals, lack of either pathway caused no major overt phenotype, although GDF15 inhibition was accompanied by increased inflammatory activation in skeletal muscle, indicating that its immunomodulatory role is not restricted to the diseased state. As the redox-supporting and anti-inflammatory actions of these metabokines can be interpreted as protective, these findings raise the possibility that either the proteins themselves or their major downstream metabolic effectors could hold therapeutic value in mitochondrial diseases that fail to induce this response, particularly forms with neurological involvement.

---

**Supplementary Figure 1. GDF15 inhibitor treatment did not cause strong changes in physiological and pathological parameters of MM mice.**

(A) Weekly measurements of mouse body weight in the span of the experiment. n = 3-8/group.

(B) Mouse food consumption on the last week of the treatments for 4 days. n = 3-8/group.

(C) Measurements of glucose levels during the glucose tolerance test (GTT). n = 3-8/group.

(D) Fold changes of mitochondrial DNA (mtDNA) copy number. n=3-6/group.

(E-F) mtDNA deletion load in mouse skeletal muscles; visualization and quantification of the PCR assay. Long range used for detection of deletions, short – for evaluation of total mtDNA load. Quantification done as a ratio of intensities between long- and short-range PCR bands. n=3/group.

(G) Differentially expressed genes in deletor muscles without inhibition of GDF15 signaling as compared to vehicle treated WT. n=4.

(H) Western blotting of ISRmt genes in skeletal muscle. n = 5/group.

(I-J) Ratio between the phosphorylated and total forms of (I) eIF2alpha and (J) S6 proteins as quantified from Fig. 1H. n=3-4.

(K-L) Fold changes of key ISRmt genes in (K) heart and (L) muscle. n=3-4/group.

Abbreviations: WT – wild type, DEL – deletor, DELFKO – deletor FGF21 knockout, VEH – vehicle, INH – inhibitor. Star significance: ns – nonsignificant, p>0.05, * - p ≤ 0.05, ** - p ≤ 0.01, *** - p ≤ 0.001, **** - p ≤ 0.0001.

**Supplementary Figure 2. GDF15 regulates expression of immune response genes**

(A) Gene expression levels of various genes involved in immune response in skeletal muscle. n=3-4.

Abbreviations: WT – wild type, DEL – deletor, DELFKO – deletor FGF21 knockout, VEH – vehicle, INH – inhibitor. Star significance: ns – nonsignificant, p>0.05, * - p ≤ 0.05.

**Supplementary Figure 3. GDF15 and FGF21 crosstalk in regulation of muscle and heart metabolism**

(A) Metabolic levels of palmitoleic acid in skeletal muscle. n=3-6.

(B) Significantly changed metabolites in muscles of WT mice lacking GDF15 signaling activity.

(C) Metabolic changes in hearts of deletors with inhibited GDF15 signaling. Comparison done between the same genotypes but different treatments.

(D) Gene expression levels of ASNS enzyme in the heart. n=3-4.

(E-F) Metabolic changes in hearts of deletors without FGF21 (E) without and (F) with GDF15 inhibition as compared to vehicle-treated controls.

(G-H) Differentially expressed genes after inhibited GDF15 activity in (G) deletors without FGF21 and

(H) wild types.

Abbreviations: WT – wild type, DEL – deletor, DELFKO – deletor FGF21 knockout, VEH – vehicle, INH – inhibitor. Star significance: ns - p>0.05, * - p ≤ 0.05, ** - p ≤ 0.01, *** - p ≤ 0.001, **** - p ≤ 0.0001.

**Supplementary Figure 4. GDF15 shows complexity in its function depending on the organ and genomic background**

(A) Fold change of significantly changed (pvalue < 0.05) mitochondrial OXPHOS subunits from RNA- Seq data of the skeletal muscle. Subunits sorted according to their complex. Circle – nuclear encoded subunits, triangle – mtDNA-encoded subunits.

(B) Expression profiles of representative genes from (S3B) – Tnni3, Rps5, Mrps16. n=3-4.

(C - E) Differentially expressed genes in hearts of (C) deletors lacking GDF15 and FGF21 signaling,

(D) WT and (E) deletors with inhibited GDF15 activity; volcano plots and pathway analyses. Comparisons done between the same genotypes but different treatments.

(F - H) Differentially expressed genes in livers of (F) deletor and (G) WT mice lacking GDF15 signaling,

(H) deletors lacking GDF15 and FGF21 activity; volcano plots and pathway analyses. Comparisons done between the same genotypes but different treatments.

## STAR★Methods

### Key resources table

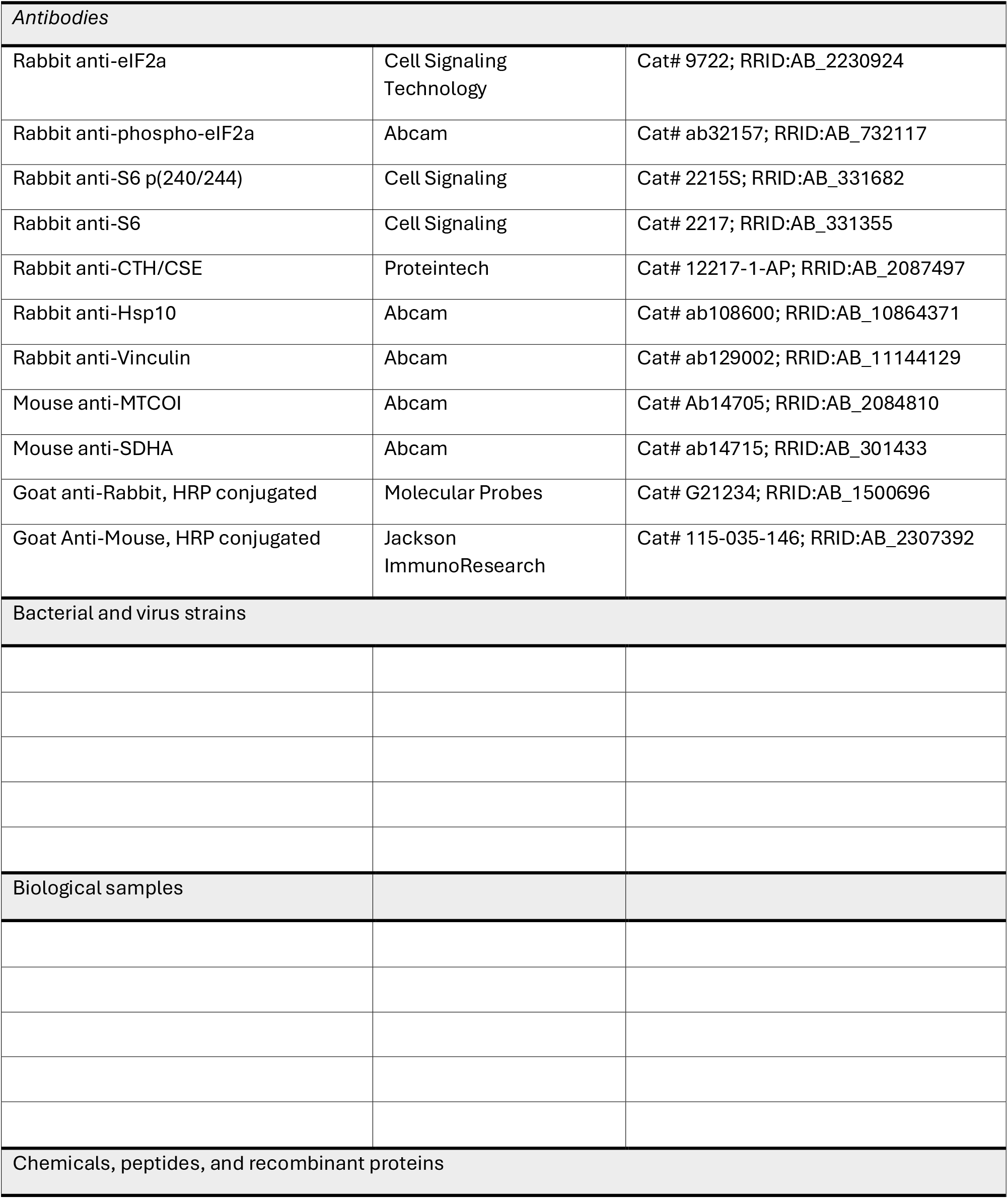
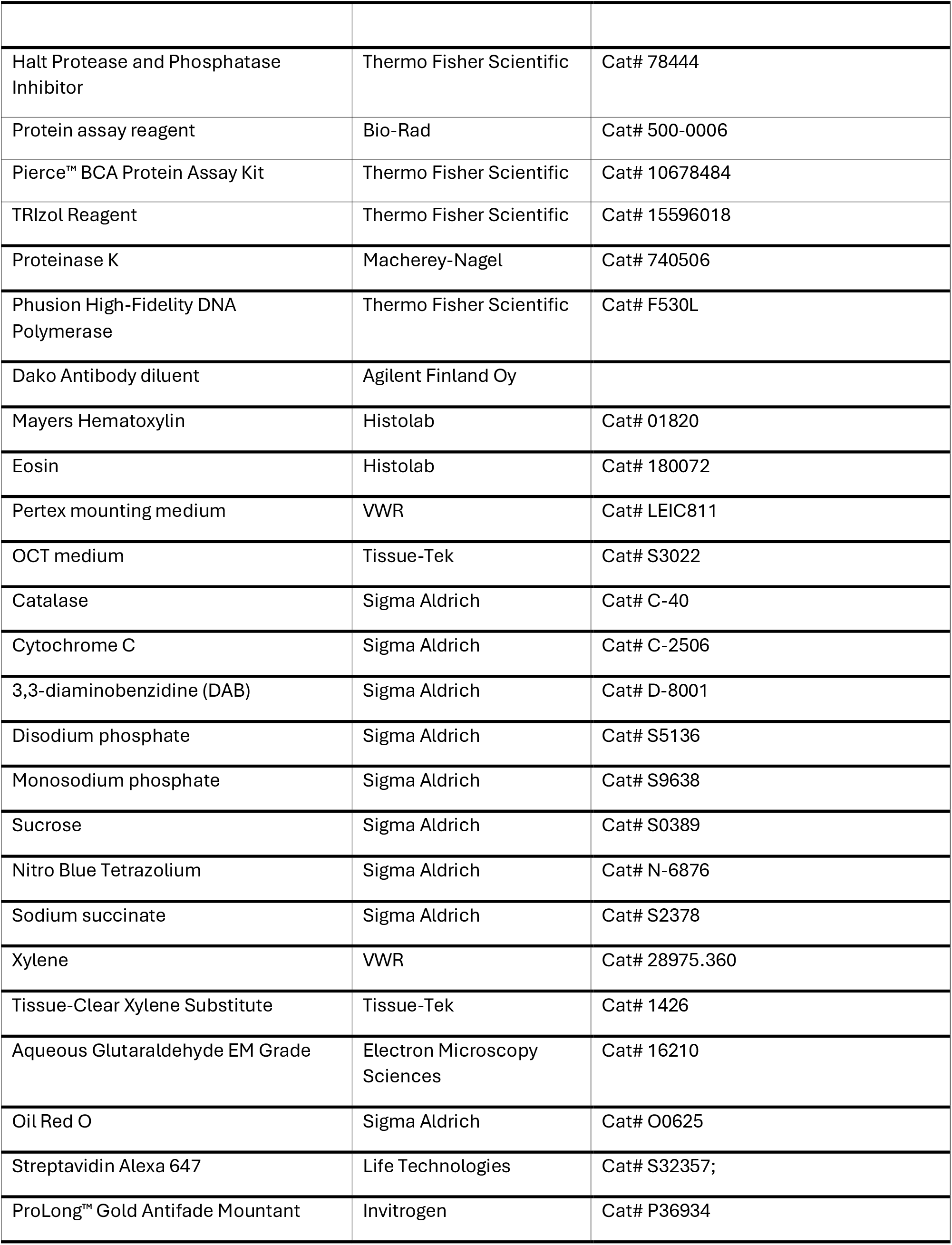
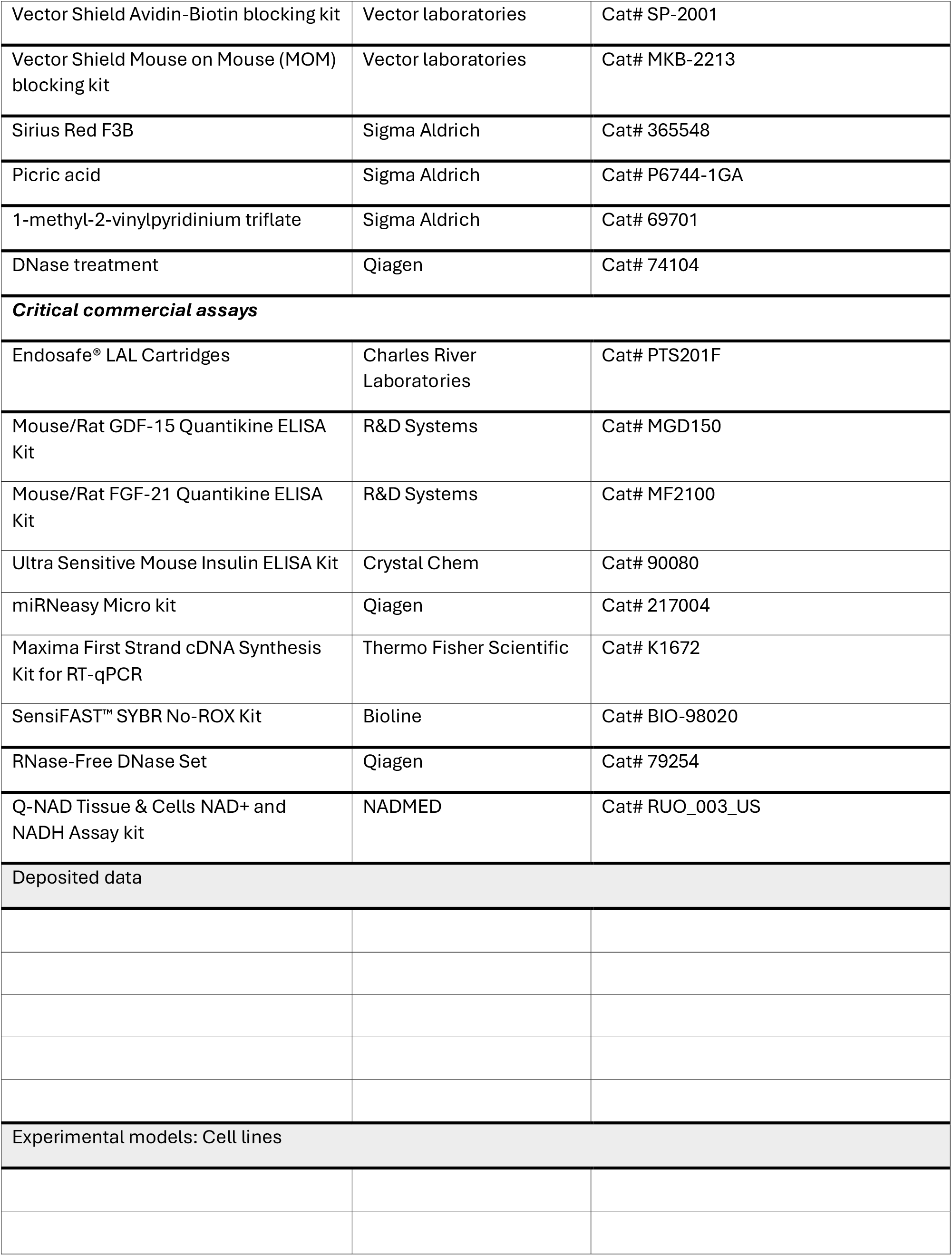
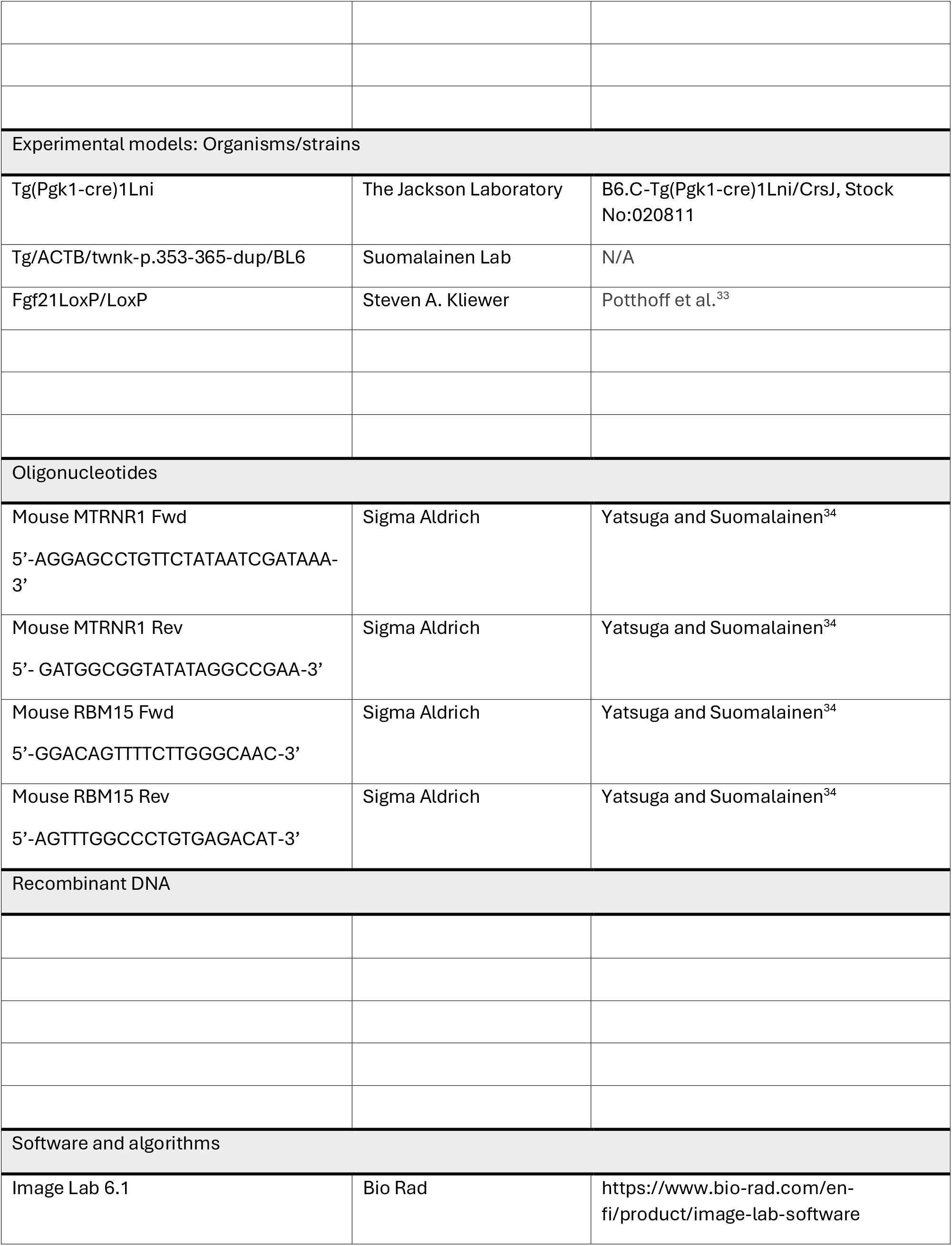
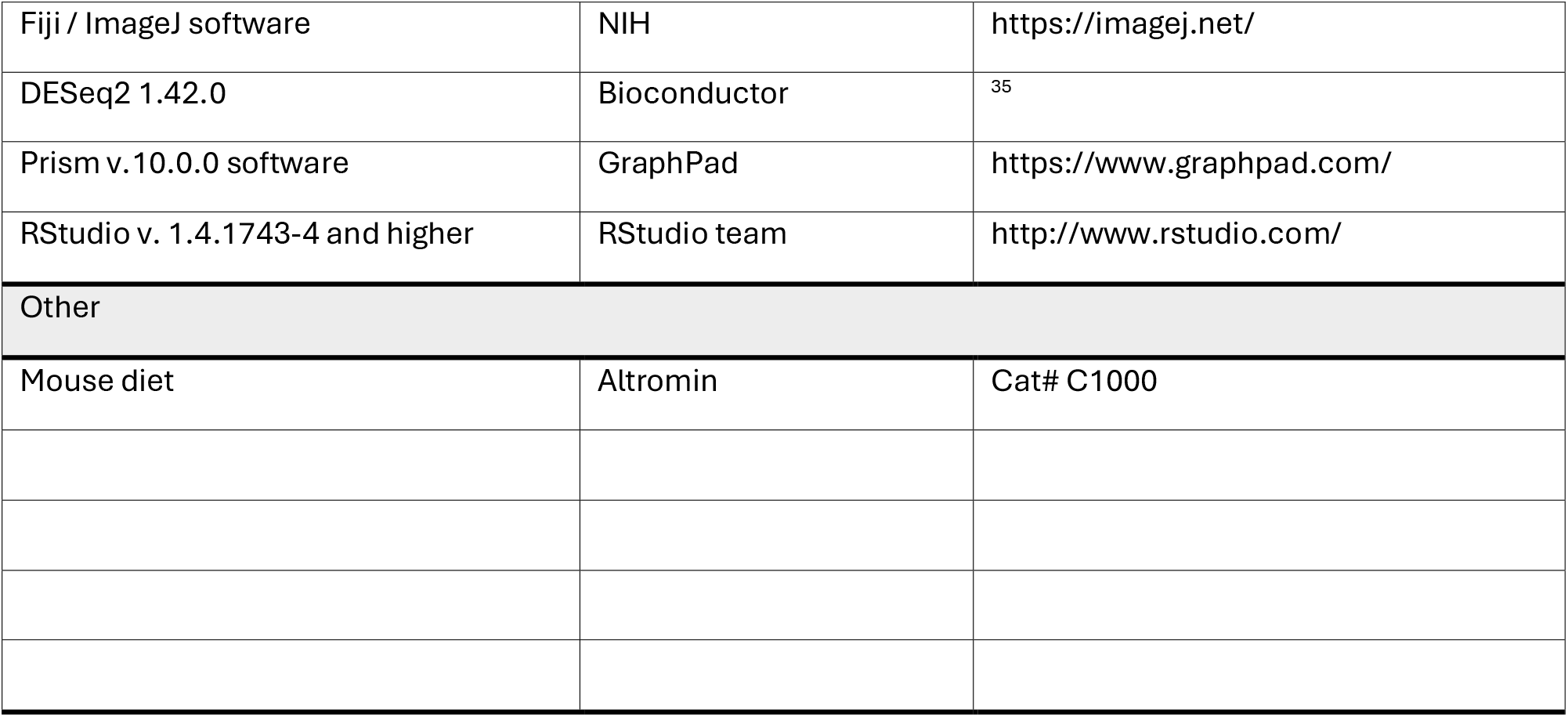

### Resource availability

#### Lead contact

Further information and requests for resources and reagents should be directed to and will be fulfilled by the lead contact, Anu Suomalainen

#### Materials availability

Requests regarding the GDF15 neutralizing antibody may be directed to NGM Biopharmaceuticals

#### Data and code availability

RNA-seq data have been deposited in the European Nucleotide Archive (ENA) under accession number PRJEB115121. Raw metabolomics mass spectrometry data generated in this study have been deposited in the Mass Spectrometry Interactive Virtual Environment (MassIVE) repository and are publicly available under accession number MSV000102200. This paper does not report original code. Any additional information required to reanalyze the data reported in this work paper is available from the lead contact upon request.

## Acknowledgements

We thank NGM Biopharmaceuticals and its members (Harding Luan, Arthur Hsu and Julie Roda) for providing the GDF15 antibody, financial support, and advice on data analysis and manuscript preparation. We gratefully acknowledge funding from Research Council of Finland for support, including the funding for MetaScale Centre of Excellence (to A.S. #361873, #345248 #374084). We also thank the technical staff of the ASW laboratory: Markus Innilä for invaluable contribution to histology; Swagat Pradhan, Takayuki Mito, and Tuula Manninen for with animal tissue collection; and Yilin Kang and Saara Forsström for feedback on the initial version of the manuscript.

## Author contributions

A.M., A.S. and representatives of NGM Biopharmaceuticals conceptualized the study. A.M. designed the experiments and performed most experimental procedures and data analysis. A.M. and A.S. wrote and finalized the manuscript; N.K. contributed to histology analysis. L.E. contributed to NAD and GSH analyses, method development and manuscript editing. S.J. and M.I. contributed to method development and analysis. All authors edited and commented the manuscript.

## Declaration of interests

A.S. and L.E. are co-founders and shareholders of NADMED Ltd, Helsinki, Finland. A.S. is the chair of the board of NADMED Ltd.

This study received financial support from NGM Biopharmaceuticals for A.M. salary and experimentation, including the GDF15-inhibiting antibody.

## Experimental model and study participant details

### Ethical Approval

All the procedures and experiments with mice were performed in accordance with national and international guidelines. Animal handling licenses were approved by the Finnish Committee of Experimental Animal research.

### Animal models

In this study, we utilized previously reported mouse models. Deletor mice overexpressing mitochondrial Twinkle helicase (*Twnk*) transgene with a dominant mutation (duplication of amino acids p.353-365) also found in patients with mitochondrial myopathy^1,7^. FGF21 knockout mice with deletor background. FGF21 deleted using the Cre-LoxP recombination system as described in Forsström and Jackson et al.^9^. Wild types were littermates with the same congenic strain, C57BL/6, as other models. Male mice were used due to the slightly stronger myopathy phenotype than in females. Animals of 22-24 months of age with active phenotype were used.

Animal genotypes abbreviations:

WT – wild type

DEL – deletor

DEL-FKO – deletor

FGF21 knockout.

Animals were housed at room temperature of 22°C with a controlled 12 hours light/dark cycle. Food and water were available *ad libitum*. Prior to sacrifice and tissue collection, mice were fasted overnight, and food was returned at 6 am on the day of collection for similar food uptake control.

## Method details

### Anti-GDF15 Antibody Generation

The variable regions of heavy chain and light chain of the GDF15 neutralizing antibody were subcloned into the pTT5 vector. The plasmids encoding the light and heavy chains were co-transfected into Expi293 cells using the ExpiFectamine Transfection kit (Thermo Fisher Scientific), and the conditioned media was harvested, clarified by centrifugation, and sterile-filtered. Antibodies were purified by protein A affinity using a HiTrap MabSelect SuRe column (Cytvia) followed by desalination and ion exchange and eluted using a 100 mL linear gradient (0–500 mM NaCl) in 25 mM sodium acetate (pH 5.5). Fractions containing aggregate-free IgG, as confirmed by SDS-PAGE, were pooled, and the final product was concentrated using a 30 kDa MWCO filter and formulated into 25 mM HEPES, 150 mM NaCl (pH 7.5). Antibody identity was confirmed by QTOF mass spectrometry (Agilent, 6520 Accurate Mass QTOF, Model G6520B). All samples were confirmed to be >98% monodispersed by size exclusion chromatography using a Yarra SEC-3000 column (Phenomenex) on an Agilent 1200 series HPLC. Endotoxin levels were < 1 EU/mg protein (tested with Endosafe® LAL Cartridges).

### Administration of GDF15-blocking antibody and mock control

The GDF15-specific and control antibodies were dissolved in PBS and administered weekly intraperitoneally for 4 weeks. Weekly dosing was used to maintain continuous exposure, based on the established pharmacokinetic properties of recombinant IgG antibodies in mice. Total of 5 injections were performed with the 5^th^ one 24 hours before sacrifice. Working concentration was 10 mg/kg of body weight.

Treatment abbreviations:

Veh - vehicle

Inh - GDF15 binding antibody

Weight was monitored twice a week throughout the entire length of the experiment. CLAMS recordings were done in week 3. Following the 4^th^ injection, food consumption was recorded. Food was weighed, placed into mouse cages, and re-weighed daily for 4 days. Food consumption was evaluated by the amount of food (in grams) consumed within 24 hours. At the end of week 4, a glucose tolerance test (GTT) was performed.

### Tissue collection

Prior to tissue collection, mice were similarly fed: food removed the previous day at 7 pm for overnight fasting and returned at 6-7 am on the day of termination to ensure all were in a similar state of feeding. Body temperature was measured just before euthanasia. Animals were sacrificed by CO2 and blood was collected by the heart puncture after which the mice were decapitated. Part of the whole blood was frozen in liquid nitrogen and the other part was left on ice to coagulate. It was later centrifuged to separate serum, which was then aliquoted and flash frozen in liquid nitrogen. Collected organs were either flash frozen for biochemical analyses, frozen for cryosections or fixed in 4% Paraformaldehyde for histology. All the frozen materials were transferred for long-term storage at -80°C.

### Respiratory Function Testing in Animals

Oxygen consumption, carbon dioxide production and activity were measured using the Oxymax Lab Animal Monitoring System (CLAMS, Columbus Instruments). Mice were monitored on the third week (after 14 days) of treatment. Animals were housed in individual cages for 48 hours. The first 24 hours were not recorded to allow for animals to adjust to the new cage conditions followed by recordings for 24 more hours. Conditions inside the cages remained constant: room temperature of 22°C and 12-hour light/dark cycle. Rates of O_2_ consumption (vO_2_) and CO_2_ production (vCO_2_) were measured by the system and normalized to animal body weight. Respiratory exchange ratio (RER) was calculated as the ratio of the vCO_2_ to vO_2_.

### Glucose Tolerance Test (GTT)

Prior to the assay, animals were placed in clean cages and starved overnight. The food was returned to the cages at 6 am, mice were re-fed for 2 hours to achieve similar food intake and starved again for 4 hours. Glucose was dissolved in PBS at concentration of 0.2mg/µl. Glucose solution was administered intraperitoneally at 2g/kg. Blood was collected from tail cuts at time points of 0, 15, 30, 90, and 120 min for glucose measurements. Extra blood samples were collected at 0, 30 and 120 min for later insulin measurements. It was immediately centrifuged at 200 xg for 5 min. Plasma was transferred to fresh tubes and flash frozen in liquid nitrogen.

### Protein analysis by western blot

Total protein was isolated from skeletal muscle. First, 25 mg of tissue was homogenized using Precellys beads (Bertin Technologies) for hard tissues in 20-30 volumes of PBS with protease and phosphatase inhibitors at 5500 rpm, 30 sec. The homogenized tissue was transferred to clean Eppendorf tubes. 5x RIPA buffer was added to reach 1x concentration. The tissue was lysed on ice for 20 min and centrifuged at 10000 xg, +4°C for 30 min to remove the insoluble debris. Clear supernatants were transferred to fresh tubes. Protein concentration was measured with Bradford assay. Samples were diluted to the desired concentration and stored in 1x Laemmli buffer.

Proteins were separated on SDS-PAGE gels (TGX 4-20% Stain Free, Biorad) and then transferred onto PVDF membrane using semidry method. The membranes were blocked for 1 hr in 5% milk in 1xTBS-T (Tris Buffered Saline with 0.1% Tween20). Primary antibodies were incubated overnight at 4°C for overnight shaking at concentrations according to manufacturer’s instructions. Secondary HRP- conjugated antibodies were added at 1:10000 dilution in 5% milk in 1xTBS-T for 1 hr incubation at room temperature. The signal was visualized by Clarity Western ECL Substrate and imaged using ChemiDoc™ XRS+ System (Biorad). Signal intensities were normalized and quantified by ImageLab 6.1.0 (Biorad).

### mtDNA Copy Number Measurements

Total cellular DNA was isolated from mouse skeletal muscle following the standard proteinase K and phenol-chloroform extraction methods.

Two sets of primers were used for mtDNA Load Evaluation: 1) primers targeting MTRNR1 gene at 12S region of mtDNA and 2) primers against a nuclear-encoded one-copy gene RBM15. Products of the reactions were run on standard 1% agarose gel. Band intensities of mitochondrial amplification products were normalized to band intensities of the nuclear target.

### mtDNA Deletion Load Analysis

MtDNA deletion load was determined by a semiquantitative long-template PCR amplification^36^, using primers flanking a large region between 16S rRNA and NDI of mtDNA to allow for detection of deleted fragments. In the reaction, 10 µg of total muscle DNA was amplified with the appropriate primers (5’- GAGGTGATGTTTTTGGTAAACAGGCGGGGT-3’ and 5’-GGTTCGTTTGTTCAACGATTAAAGTCCTACGTG-3’),

Phusion DNA polymerase and GC buffer (Thermo Fisher). Long-template PCR program included the initial denaturation step at 98°C for 30 sec, 22 cycles at 98°C for 10 sec, extension at 72°C for 12 min, and the final step at 72°C for 10 min. Total undeleted mtDNA amount was also measured using 10 µg DNA, Phusion polymerase and GC buffer but a different set of primers (5’- ACCCCGCCTGTTTACCAAAAACATCACCTC-3’ and 5’-ACGTACCACTTTAATCGTTGAACAAACGAACC-3‘). All the PCR products were run on a standard 1% agarose gel, visualised by ChemiDoc™ XRS+ System (Bio- Rad), and the deleted products were compared to the total mtDNA amplicons.

### Histology

#### Respiratory Chain Enzyme Activity and Oil-red-O, Fresh-frozen Tissues

Tissues were frozen by embedding in OCT medium and placing into a 2-methylbutane bath cooled by liquid nitrogen.

Simultaneous histochemical activity analysis of mitochondrial cytochrome c oxidase (COX) and succinate dehydrogenase (SDH) was done in mouse skeletal muscles. The tissues were incubated for 30 min at room temperature in COX incubation buffer consisting of 0.05 M phosphate buffer (pH 7.4), 3, 3 – diaminobenzidine (DAB), catalase, cytochrome c and sucrose. SDH treatment was for 40 min at 37°C with the buffer formulation of 0.05 M phosphate buffer (pH 7.4) with nitro-blue tetrazolium and sodium succinate. After the reactions, tissue slices were dehydrated in ascending alcohols followed by xylene treatment and mounting. Quantification of COX negative, SDH positive, and total fiber numbers was done in Fiji (Image J) software.

Frozen livers were used for lipid droplet analysis by Oil-Red-O stainings. For the staining, tissue sections were fixed in formic calcium followed by 10 min incubation in Oil-Red-O solution. Brief staining with hematoxylin was included to visualize the nuclei.

#### CD68 and CD45 immunohistochemistry, Fresh-frozen Tissues

For immunohistochemical detection of CD68 and CD45, frozen tissue sections were air-dried at room temperature for 2–4 h, fixed in ice-cold acetone for 5 min, and washed in PBS. Endogenous peroxidase activity was quenched with freshly prepared methanol/H₂O₂, followed by washes in water and PBS containing 0.1% Tween-20 (PBS-T). Sections were then outlined with a PAP pen, blocked for 1 h at room temperature in freshly prepared 7.6% normal horse serum in PBS-T, and incubated overnight at 4°C in a humid chamber with primary antibodies diluted in DAKO REAL antibody diluent (anti-CD68, 1:600; anti- CD45, 1:50), while parallel control sections received rat IgG at matched concentrations. On the following day, sections were washed in PBS-T, incubated for 30 min at room temperature with ImmPRESS goat anti-rat secondary polymer reagent, and developed with freshly prepared DAB solution activated immediately before use with H₂O₂. After additional washes, sections were counterstained with Mayer’s hematoxylin for 30 s, rinsed in running tap water, washed in RO water, dehydrated through graded ethanol and xylene, and mounted with Pertex.

#### COX/SDH staining in cortex from paraffin fixed tissues

In paraffin sections, COX and SDH immunodetections were performed on separate slides. First, the sections were deparaffinized in descending alcohols. For antigen retrieval, the samples were pre-treated with 10 mM citrate buffer in 1 mM EDTA (pH 6.0) and incubated in a decloaking pressure cooker at 110°C for 1 hr. After washes in PBS-Tween buffer, endogenous peroxidase activity was quenched by incubation in 1% methanol solution in H_2_O_2_ followed by washes and blocking in serums (5% Fetal Bovine serum + 7.5% Horse normal serum in 0.1% Tween-PBS) for 1 hr at room temperature. Primary antibodies (mtCo1; SDHA) and mouse IgG negative control were diluted in DAKO diluent to 2.5 µg/mL and incubated overnight at 4°C. Detection of primary antibodies was done using Avidin-biotin system by incubation with avidin-biotin-peroxidase complex for 30 min at room temperature. After brief immersion in hematoxylin solution, the tissues were dehydrated in ascending alcohols and fixed in mounting media.

#### Sirius Red staining for collagen paraffin fixed tissues

Paraffin sections were dewaxed and hydrated. The nuclei were stained with haematoxylin for 8 minutes, and then the slides were washed for 10 minutes in running tap water followed by incubation in picro- sirius red (0.5 gr Sirius Red in 500 mL picric acid) for 1 hr. After that, the slides were washed twice in acidified water (5 mL glacial acetic acid to 1 L of water). The water was then removed by vigorous shaking, and the samples were rehydrated in three changes of 100% ethanol. Finally, the slides were cleared in xylene and mounted in a resinous medium.

### RNA isolation and sequencing

RNA isolation was done using a RNeasy micro kit with DNase treatment following the manufacturer’s instructions. Quality of the purified RNA was evaluated by TapeStation RNA QC assay. RNA-Sequencing was done with the “Bulkseq” method of counting 3’UTR for gene expression. Data was provided as gene names and individual counts. Both RNA quality control and sequencing were made at the Functional Genomics Unit (FuGu), University of Helsinki, Finland.

Data analysis was done using a broadly used R-package DEseq2 and pathway analysis - in Metascape online resource.

### Targeted Metabolomics

Targeted metabolomics was performed at the local metabolomics unit (FIMM META) from frozen muscle and heart tissues. Method: Orbitrap Full Profiling of 462 metabolites. Metabolite extraction was done in 40:40:20 ACN:Methanol:Water. Samples were normalized with normalizing factor counted based on weight. Data analysis was done using the conventional Metaboanalyst platform.

### NAD^+^/NADH, NADP^+^/NADPH and GSH/GSSG measurements

Quantitative analysis of reduced and oxidized glutathione and NAD/NADP metabolites in frozen tissue samples was done in accordance with the manufacturer’s instructions of the kit or as a laboratory service by NADMED Ltd.

For normalization of the results, protein content in the sample was measured using the pellet obtained after centrifugation of the homogenate. For protein assay BCA kit was used.

### Quantification and statistical analysis

Initial data analyses (in vivo observations, histology, biochemical methods, RNA-Seq) were performed blindly. GDF15 inhibitor and vehicle were labelled as vial 1 and 2 respectively.

GraphPad Prism (versions 10.0.0 and above; GraphPad Software, USA) was used for statistical analyses and graphical presentation of the data. Statistical analysis for multiple groups was performed as a two-way ANOVA with multiple comparisons according to Sidak method. In bar plots, error bars represent standard deviation (SD) whilst in box plots, error bars are the minimum and maximum values. The center bar of box plots represents median. P-values of less than 0.05 were considered statistically significant.

RNA-Seq data was analyzed using DESeq2 package^35^ with volcano plots data visualization done in R (ver 4.1.3 and above). Volcano plots were made using Enhanced Volcano package^37^.

Metabolomics data was partially visualized in R for volcano plots. Samples were normalized with normalizing factor counted based on weight. Main data analysis for metabolomics was performed in MetaboAnalyst 6.0 online software^38^. Pathway analysis was performed in Metascape^39^.

Analysis and quantification of histological samples were performed in ImageJ software (NIH; Bethesda, MD, USA).

