## Supplementary figures for "GDF15 modifies NAD+ salvage pathway and dampens immune response in mitochondrial disease"

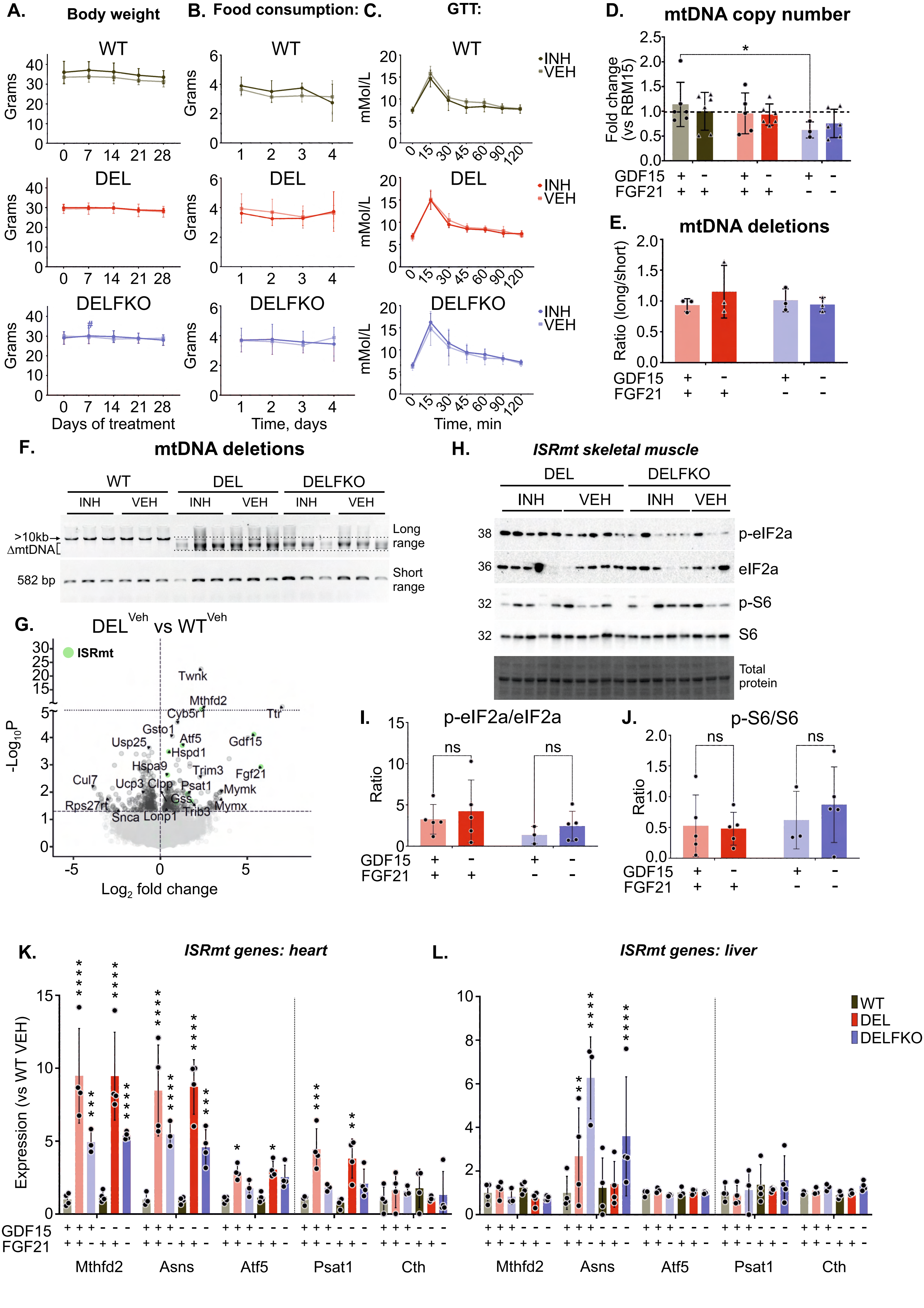
A. Body weight
B. Food consumption:
C. GTT:
WT
DEL
DELFKO
Grams
Days of treatment
Time, days
mMol/L
Time, min
INH
VEH
D. mtDNA copy number
Fold change (vs RBM15)
GDF15
FGF21
E. mtDNA deletions
Ratio (long/short)
F. mtDNA deletions
>10kb
ΔmtDNA
582 bp
Long range
Short range
G. DEL^Veh vs WT^Veh
ISRmt
-Log10P
Log2 fold change
Twnk
Mthfd2
Cyb5r1
Ttr
Gsto1
Atf5
Usp25
Hspd1
Gdf15
Hspa9
Trim3
Fgf21
Cul7
Ucp3
Clpp
Psat1
Mymk
Gss
Rps27rt
Mymx
Snca
Lonp1
Trib3
H. ISRmt skeletal muscle
p-eIF2a
eIF2a
p-S6
S6
Total protein
I. p-eIF2a/eIF2a
Ratio
ns
J. p-S6/S6
K. ISRmt genes: heart
Expression (vs WT VEH)
Mthfd2
Asns
Atf5
Psat1
Cth
L. ISRmt genes: liver
WT
DEL
DELFKO

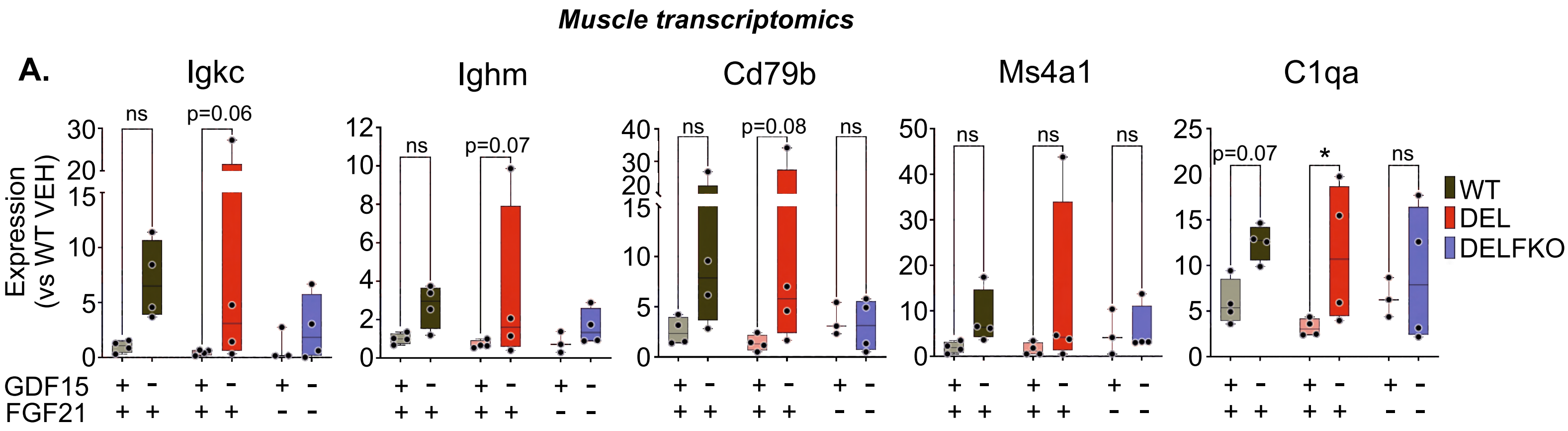
Muscle transcriptomics
A.
Igkc
Ighm
Cd79b
Ms4a1
C1qa
Expression
(vs WT VEH)
ns
p=0.06
p=0.07
p=0.08
*
GDF15
FGF21
WT
DEL
DELFKO

Figure S2

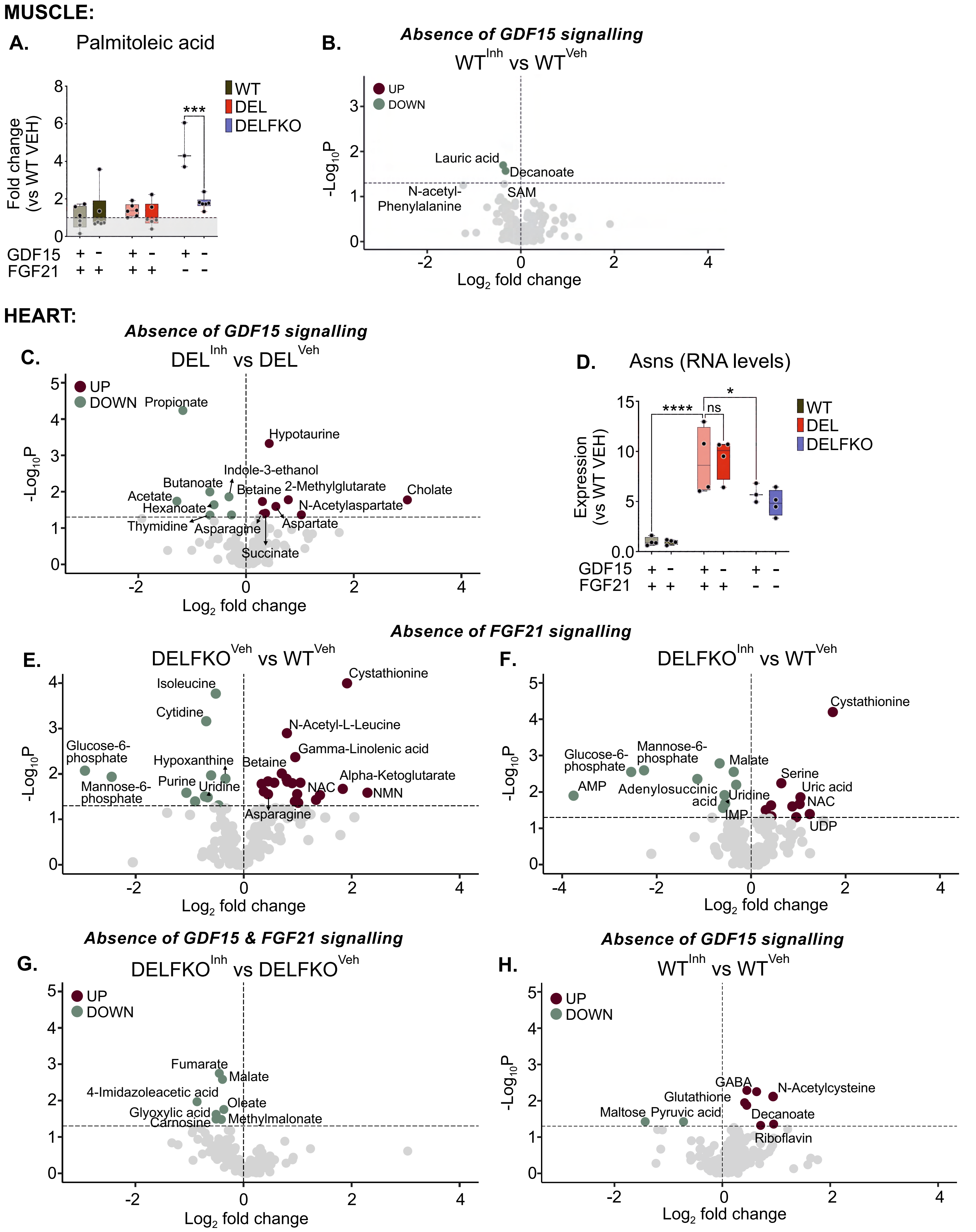
MUSCLE:
A.
Palmitoleic acid
Fold change (vs WT VEH)
WT
DEL
DELFKO
***
GDF15 + - + - + -
FGF21 + + + + - -
B.
Absence of GDF15 signalling
WT^Inh vs WT^Veh
UP
DOWN
Lauric acid
Decanoate
N-acetyl-Phenylalanine
SAM
-Log10P
Log2 fold change
HEART:
Absence of GDF15 signalling
C.
DEL^Inh vs DEL^Veh
UP
DOWN
Propionate
Hypotaurine
Indole-3-ethanol
Butanoate
Betaine
2-Methylglutarate
Cholate
Acetate
Hexanoate
N-Acetylaspartate
Thymidine
Asparagine
Aspartate
Succinate
-Log10P
Log2 fold change
D.
Asns (RNA levels)
Expression (vs WT VEH)
****
ns
*
WT
DEL
DELFKO
GDF15 + - + - + -
FGF21 + + + + - -
Absence of FGF21 signalling
E.
DELFKO^Veh vs WT^Veh
Isoleucine
Cystathionine
Cytidine
N-Acetyl-L-Leucine
Gamma-Linolenic acid
Glucose-6-phosphate
Hypoxanthine
Betaine
Alpha-Ketoglutarate
Mannose-6-phosphate
Purine
Uridine
NAC
NMN
Asparagine
-Log10P
Log2 fold change
F.
DELFKO^Inh vs WT^Veh
Cystathionine
Glucose-6-phosphate
Mannose-6-phosphate
Malate
AMP
Adenylosuccinic acid
Serine
Uric acid
Uridine
NAC
IMP
UDP
-Log10P
Log2 fold change
Absence of GDF15 & FGF21 signalling
G.
DELFKO^Inh vs DELFKO^Veh
UP
DOWN
Fumarate
Malate
4-Imidazoleacetic acid
Oleate
Glyoxylic acid
Carnosine
Methylmalonate
Log2 fold change
Absence of GDF15 signalling
H.
WT^Inh vs WT^Veh
UP
DOWN
GABA
N-Acetylcysteine
Glutathione
Maltose
Pyruvic acid
Decanoate
Riboflavin
-Log10P
Log2 fold change

**Figure S3**

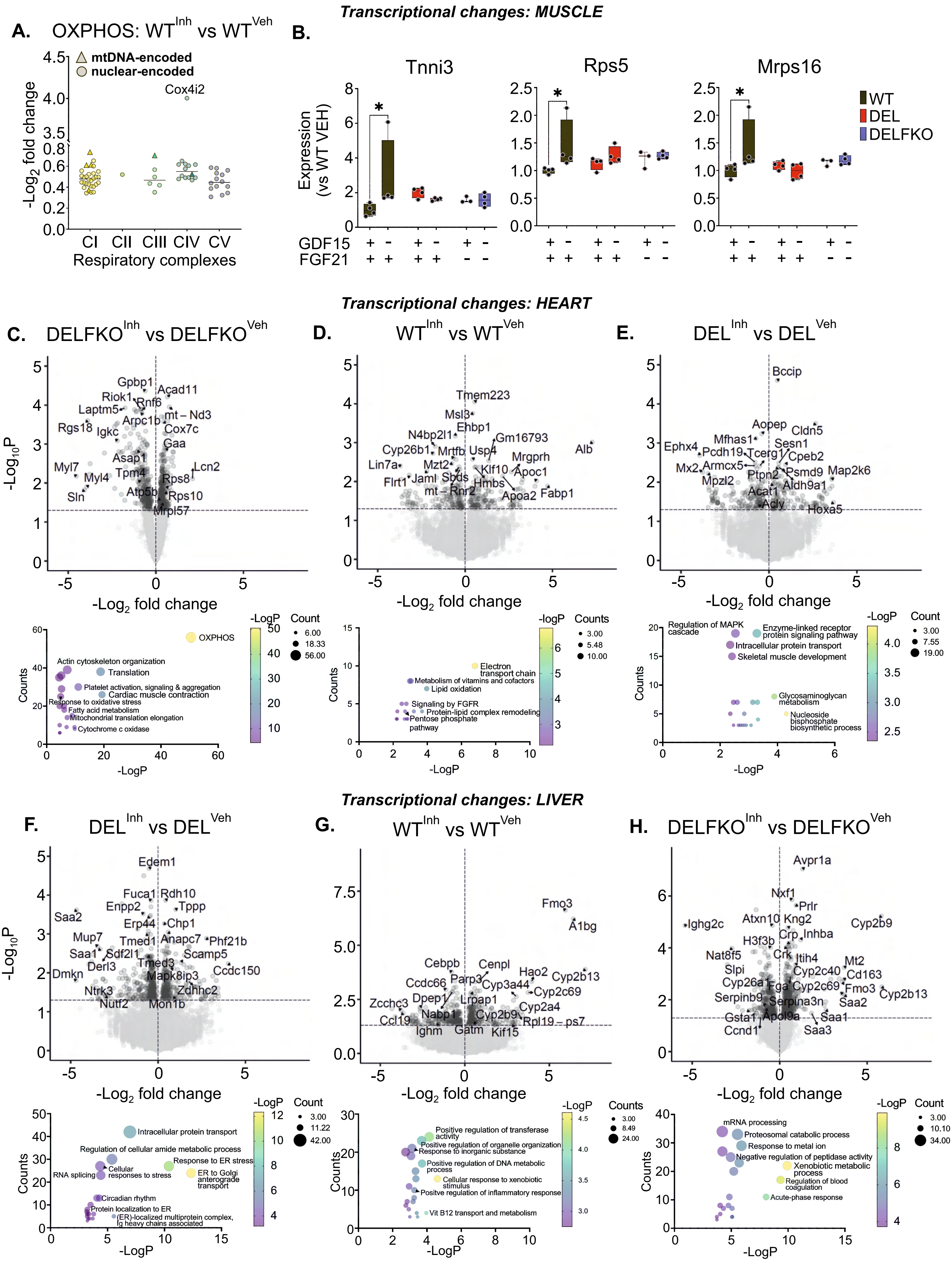
Transcriptional changes: MUSCLE
A. OXPHOS: WTInh vs WTVeh
mtDNA-encoded
nuclear-encoded
Cox4i2
-Log2 fold change
CI CII CIII CIV CV
Respiratory complexes
B.
Tnni3
Rps5
Mrps16
Expression (vs WT VEH)
WT
DEL
DELFKO
GDF15
FGF21
Transcriptional changes: HEART
C. DELFKOInh vs DELFKOVeh
Gpbp1
Riok1
Acad11
Rnf6
Laptm5
mt – Nd3
Arpc1b
Rgs18
Igkc
Cox7c
Gaa
Asap1
Lcn2
Myl7
Tpm4
Myl4
Rps8
Atp5b
Rps10
Sln
Mrpl57
-Log10P
-Log2 fold change
OXPHOS
Actin cytoskeleton organization
Translation
Platelet activation, signaling & aggregation
Cardiac muscle contraction
Response to oxidative stress
Fatty acid metabolism
Mitochondrial translation elongation
Cytochrome c oxidase
Counts
-LogP
Count
D. WTInh vs WTVeh
Tmem223
Msl3
Ehbp1
N4bp2l1
Gm16793
Cyp26b1
Mrtfb
Usp4
Alb
Mrgprh
Lin7a
Mzt2
Klf10
Apoc1
Sbds
Flrt1
Jaml
Hmbs
mt – Rnr2
Apoa2
Fabp1
Electron transport chain
Metabolism of vitamins and cofactors
Lipid oxidation
Signaling by FGFR
Protein-lipid complex remodeling
Pentose phosphate pathway
-logP
Counts
E. DELInh vs DELVeh
Bccip
Aopep
Cldn5
Mfhas1
Sesn1
Ephx4
Pcdh19
Tcerg1
Cpeb2
Mx2
Armcx5
Psmd9
Map2k6
Ptpn2
Mpzl2
Aldh9a1
Acat1
Acly
Hoxa5
Regulation of MAPK cascade
Enzyme-linked receptor protein signaling pathway
Intracellular protein transport
Skeletal muscle development
Glycosaminoglycan metabolism
Nucleoside bisphosphate biosynthetic process
Transcriptional changes: LIVER
F. DELInh vs DELVeh
Edem1
Fuca1
Rdh10
Enpp2
Tppp
Saa2
Erp44
Chp1
Mup7
Tmed1
Anapc7
Phf21b
Saa1
Sdf2l1
Scamp5
Derl3
Tmed3
Ccdc150
Dmkn
Mapk8ip3
Ntrk3
Zdhhc2
Nutf2
Mon1b
Intracellular protein transport
Regulation of cellular amide metabolic process
Response to ER stress
Cellular responses to stress
RNA splicing
ER to Golgi anterograde transport
Circadian rhythm
Protein localization to ER
(ER)-localized multiprotein complex, Ig heavy chains associated
G. WTInh vs WTVeh
Fmo3
A1bg
Cebpb
Cenpl
Hao2
Cyp2b13
Ccdc66
Parp3
Cyp3a44
Cyp2c69
Zcchc3
Dpep1
Lrpap1
Cyp2a4
Ccl19
Nabp1
Cyp2b9
Rpl19 – ps7
Ighm
Gatm
Kif15
Positive regulation of transferase activity
Positive regulation of organelle organization
Response to inorganic substance
Positive regulation of DNA metabolic process
Cellular response to xenobiotic stimulus
Positive regulation of inflammatory response
Vit B12 transport and metabolism
H. DELFKOInh vs DELFKOVeh
Avpr1a
Nxf1
Prlr
Ighg2c
Atxn10
Kng2
Cyp2b9
H3f3b
Crp
Inhba
Nat8f5
Crk
Itih4
Slpi
Mt2
Cyp2c40
Cd163
Cyp26a1
Fga
Cyp2c69
Fmo3
Cyp2b13
Serpinb9
Serpina3n
Saa2
Gsta1
Apol9a
Saa1
Ccnd1
Saa3
mRNA processing
Proteosomal catabolic process
Response to metal ion
Negative regulation of peptidase activity
Xenobiotic metabolic process
Regulation of blood coagulation
Acute-phase response

**Figure S4**
